# Heat Inactivation of Aqueous Viable Norovirus and MS2 Bacteriophage

**DOI:** 10.1101/2023.08.22.554333

**Authors:** Marlee Shaffer, Kimberly Huynh, Verónica Costantini, Kyle Bibby, Jan Vinjé

## Abstract

Human norovirus is a leading cause of acute gastroenteritis often associated with contaminated food or water exposure. Many studies have used morphologically similar viruses, such as MS2 bacteriophage, and molecular detection methods to study the environmental fate and inactivation characteristics, given the historical challenges to culture human norovirus. In this study, we used human intestinal enteroids (HIEs) to analyze the heat inactivation kinetics of viable human norovirus compared to the surrogate MS2 bacteriophage. Norovirus decay rates were 0.22 min^-1^, 0.68 min^-1^, and 1.11 min^-1^ for 50°C, 60°C, and 70°C, respectively, and MS2 bacteriophage decay rates were 0.0065 min^-1^, 0.045 min^-1^, and 0.16 min^-1^ for 50°C, 60°C, and 70°C, respectively. Norovirus and MS2 bacteriophage had significantly different decay rates at all tested temperatures (p = 0.002 – 0.007). No decrease of RNA titers as measured by realtime RT-PCR for both human norovirus and MS2 bacteriophage over time was observed, indicating molecular methods do not accurately depict viable human norovirus after heat inactivation. Overall, our data demonstrate that MS2 bacteriophage is a conservative surrogate to measure heat inactivation and potentially overestimates the infectious risk of norovirus.

**IMPORTANCE:** Norovirus is the leading cause of epidemic and endemic acute gastroenteritis worldwide. Treatments to inactivate norovirus are critical to reducing the risk associated with contaminated food and water. The recent developments to replicate human norovirus in human intestinal enteroids (HIE) enables the evaluation of heat inactivation kinetics of viable norovirus. Historically, cultivable surrogate viruses, such as bacteriophage MS2, have been used to measure the environmental fate of human norovirus. Our findings indicate that compared to human norovirus, MS2 bacteriophage is a conservative surrogate to measure the effect of heat inactivation. Furthermore, this study corroborates that measuring viral RNA titers, as evaluated by PCR methods, does not correlate with persistence of viable norovirus.

## INTRODUCTION

Human noroviruses are icosahedral, non-enveloped, single-stranded RNA viruses of the family *Caliciviridae*.^1,2^ Norovirus is transmitted through the fecal-oral route via contaminated water and food, and person-to-person contact. Recent estimates demonstrate that noroviruses are the most common cause of acute gastroenteritis among all age groups, worldwide resulting in 685 million cases annually.^3,5^ Noroviruses cause 58% of foodborne illnesses in the United States each year with an estimated cost of $2 billion annually.^1,3^ Norovirus outbreaks linked to contaminated recreational, agricultural, and drinking waters have been reported throughout the past decade.^4^ Symptomatic norovirus infections are typically relatively mild for healthy individuals; however, clinical symptoms can be more severe and require hospitalization for the elderly, young children, and immunocompromised persons. ^3^

Due to the lack of a robust cell culture system for human norovirus, the effect of inactivation experiments have typically been measured by molecular methods, by cultivable surrogate viruses, or human challenge studies. ^4,5^ Molecular methods include polymerase chain reaction (PCR), which is unable to discern between viable and non-viable virions, leading to overestimation of viable viruses after treatments.^6,7^ Surrogate viruses, such as murine norovirus, feline calicivirus, Tulane virus, and MS2 bacteriophage, are morphologically similar to human norovirus. However, there is an ongoing debate on the relevance of surrogate viruses and viable human norovirus. Direct comparisons have yet to be made due to the lack of a culture system for human norovirus.^6,7^

Human intestinal enteroids (HIEs) have successfully shown replication of human norovirus indicating the presence of viable norovirus.^5,8^ HIEs recapitulate the complex environment of the gastrointestinal tract. ^5,9–11^ In this study, we compared the effect of heat on viable norovirus assessed by HIEs, and inactivation of cultivable surrogate MS2 bacteriophage. Viral RNA titers were also determined for both viruses.

## MATERIALS AND METHODS

### MS2 Bacteriophage Heat Inactivation

*Escherichia coli bacteriophage MS2* (ATCC® 15597-B1™) was cultured and recovered from *Escherichia coli* C-3000 (ATCC® 15597™), as described previously.^12^ After filtration the phage was stored at 4°C until used for the heat inactivation experiments. The double layer agar method was used to determine the titer of the MS2 bacteriophage stock.^13^

Each 2 mL sample tube had 900 μL of ultrapure molecular water and 100 μL of MS2 bacteriophage stock (6x10^9^ PFU/ml). A heat block was used to achieve the desired temperatures (50°C, 60°C, or 70°C), and the temperature was continually monitored in a blank tube of ultrapure water to ensure consistent heating throughout the experiment. At each time point, a tube was removed from the heat block and immediately added to an ice bath, and subsequently cultured using the double agar technique as previously described.^13^ 200 μL from each sample tube was aliquoted for downstream extractions and quantification and frozen at -80°C until used. We ran all experiments in duplicate with three technical replicates for each time point. All experiments used ultrapure molecular water as a negative control.

We extracted nucleic acids from each 200 μL aliquot using the AllPrep PowerViral DNA/RNA kit (Qiagen, Hilden, Germany) using Glass PowerBead tubes included with the kit. Solution PM1 was heated to 55°C, and 600 μL was added to each PowerBead tube with 6 μL of ß-mecaptoethanol (MP Biomedicals, Irvine, CA, USA). Each tube was vortexed and homogenized on a FastPrep 24 Bead Beating Instrument for 20 seconds at 4.5 M/s for four rounds. PowerBead tubes were centrifuged at 13,000g for 1 minute, and 700 μL of supernatant was transferred to a clean 2-mL microcentrifuge tube. The remaining steps followed the Qiagen protocol, and 100 μL of RNase-free water was used to elute the nucleic acids. Negative controls of ultrapure water were also extracted to ensure no contamination occurred during the extractions.

Extracted MS2 Bacteriophage RNA was quantified using reverse transcription droplet digital polymerase chain reaction (RT-ddPCR) performed on the BioRad QX2000 Droplet Digital PCR System with thermal cycling on the C1000 Touch Thermal Cycler (BioRad, Hercules, CA, USA). RNA reverse transcription and PCR amplification were performed in a single reaction using the One-Step RT-ddPCR Advanced Kit for Probes (BioRad, Hercules, CA, USA) per the manufacturer’s instructions. Oligonucleotide primer and probe sequences and thermocycling conditions are summarized in Table S1. Each RT-ddPCR plate included negative controls from the experiments, positive controls, and no-template controls of molecular water. We used QuantaSoft Version 1.7.4 (BioRad, Hercules, CA, USA) to manually threshold the RNA copy number for each reaction.

### Norovirus Heat Inactivation

A 10% stool suspension was prepared by adding 0.5 g of GII.4 Sydney[P31] positive stool to 4.5 ml of PBS. The stool suspension was vortexed for 30 seconds, kept at room temperature for 5 min, and vortexed again. The sample was sonicated for 10 sec, and solids were removed by centrifugation for 10 min at 10,000 x g. The supernatant was sequentially filtered through 5 μm, 1 μm, 0.45 μm, and 0.22 μm filters.^5^ The resulting 10% stool filtrate was aliquoted and stored at -70°C until tested.

Adult secretor-positive jejunal HIE cultures (J2 cell line) were grown at 37°C and 5% CO_2_ as undifferentiated 3D cultures as described previously with minor modifications.^5,14^ Briefly, HIEs were recovered from liquid nitrogen, suspended in 20 μl of Matrigel™ (Corning), plated in 24 well plates, and grown as 3D cultures in 500 μL IntestiCult™ (INT) Organoid Growth Medium (Stem Cell Technologies) supplemented with 10 μM Y-27632 (Sigma-Aldrich).

Duplicate 96 well plate monolayers were prepared as previously described.^5^ Briefly, HIE cultures were dissociated into a single-cell suspension in 100 μl of INT medium supplemented with 10 μM Y-27632 (Sigma-Aldrich) and plated as undifferentiated monolayers in collagen IV (Sigma-Aldrich) pre-coated 96 well plates. After 24 hours, we replaced the INT medium with a differentiation medium to induce cell differentiation. Differentiation media was prepared by mixing equal volumes of IntestiCult™ (INT) Organoid Basal Medium (Stem Cell Technologies) and complete media without growth factors (CMGF-; Advanced DMEM/F12 medium supplemented with 1x Glutamax, 10 mM HEPES, and 100 U/ml penicillin-streptomycin). Monolayers differentiated for four days at 37°C and 5% CO_2_, and we refreshed differentiation media every other day, ensuring 100% confluence before use.

Inactivation kinetics for viable human norovirus were determined at 50°C, 60°C, and 70°C. Two experiments were conducted per temperature. In each sample tube, 800 μL of infection media [CMGF-supplemented with 500 μM glycochenodeoxycholic acid (GCDCA) and 50 μM ceramide (C2)] was added to diluted 10% stool suspension. Samples were placed on a heat block at the desired temperature, and at each time point, a sample tube was removed for HIE infections. Negative controls (no virus) and treatment (no heat) controls were included.

All infections were performed in triplicate wells on 100% confluent 4-day-old differentiated HIE monolayers for each sample collected from each microcosm. Spiked aliquots consisted of infection media spiked with norovirus suspensions and non-spiked aliquots were solely infection media. Duplicate 96-well plates were inoculated with 100 µl of spiked and non-spiked aliquots that were pre-diluted in infection media. After a 1-hour incubation at 37°C and 5% CO_2_, HIE monolayers were washed twice with CMGF-and 100 μL differentiation medium containing 500 µM GCDCA and 50 µM C2 was added. For each set of infections, one plate was frozen immediately at -70°C, and a duplicate plate was incubated at 37°C, 5% CO_2_, for 72 hours and then frozen at -70°C.

Viral RNA from spiked and non-spiked aliquots, cells, and media at one hour and 72 hours after infection was extracted using the MagMax-96 Viral RNA Isolation Kit (Applied Biosystems, Foster City, CA) according to the manufacturer’s instructions. Norovirus genomic copies were quantified by real-time reverse-transcription quantitative PCR as described previously.^15^ The primer and probe sequences are shown in Table S1.^15^ Each reaction was prepared as 22 μL volume consisting of 12.50 μL 2X RT-PCR Buffer, 1.15 μL of primer and probe mixture, 1.00 μL of 25X RT-PCR Mix, 7.35 μL of molecular grade water, and 3 μL of nucleic acid extract. Three technical replicates of each nucleic acid extraction were performed, and each run had an MS2 bacteriophage control to confirm that no contamination was present. A standard curve using 10-fold serial dilutions of quantified GII.4 Sydney RNA transcripts was generated, and norovirus genomic copies from each sample were extrapolated from the curve. The real-time RT-qPCR limit of detection was 26.8 RNA copies per 1 µL (or 2.68’ 10^3^ RNA copies/well). Measurements below the limit of detection were excluded unless all samples were below the limit of detection, in which case samples were arbitrarily assigned to half of the limit of detection (14.3 RNA copies per 1 µL of RNA or 1.43’ 10^3^ RNA copies/well).

#### Statistical Analysis

All statistical analyses were completed R Studio 2022.07.2. The data shows all experimental runs and replicates’ mean and standard deviation. Significant differences in genomic copies and plaque-forming units (PFU) were analyzed using a Student’s t-test with multiple comparisons. Viable human norovirus and MS2 bacteriophage decay were analyzed using a monophasic decay model, which assumes first-order decay (*Equation 1*).

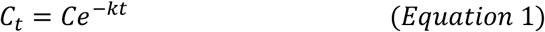

Where: *C*_*t*_ is the genome copies or PFU at time *t, C*_*0*_ is the genome copies or PFU at time 0, *k* is the decay rate, and *t* is the time in minutes.

## RESULTS

### Heat Inactivation of MS2 Bacteriophage

We calculated the monophasic decay of MS2 bacteriophage for each point using *Equation 1* **(Figure 1A**). The decay rate increased with temperature for viable MS2 bacteriophage; however, the molecular data showed no significant change in RNA signal with temperature (*p* = 0.12 – 0.93). The calculated decay rates for MS2 bacteriophage were 0.0065 min^-1^, 0.045 min^-1^, and 0.16 min^-1^ for 50°C, 60°C, and 70°C, respectively. Summaries of the decay rates, half-life, and T_90_ are shown in **Table 1**, and all viable decay rates were statistically different (p = 0.01 -<0.0001). The decay rates found for each temperature were plotted against temperature **(Figure 2**); for every degree increase in temperature (°C), the decay rate increased by 0.008 min^-1^.

**Table 1:**
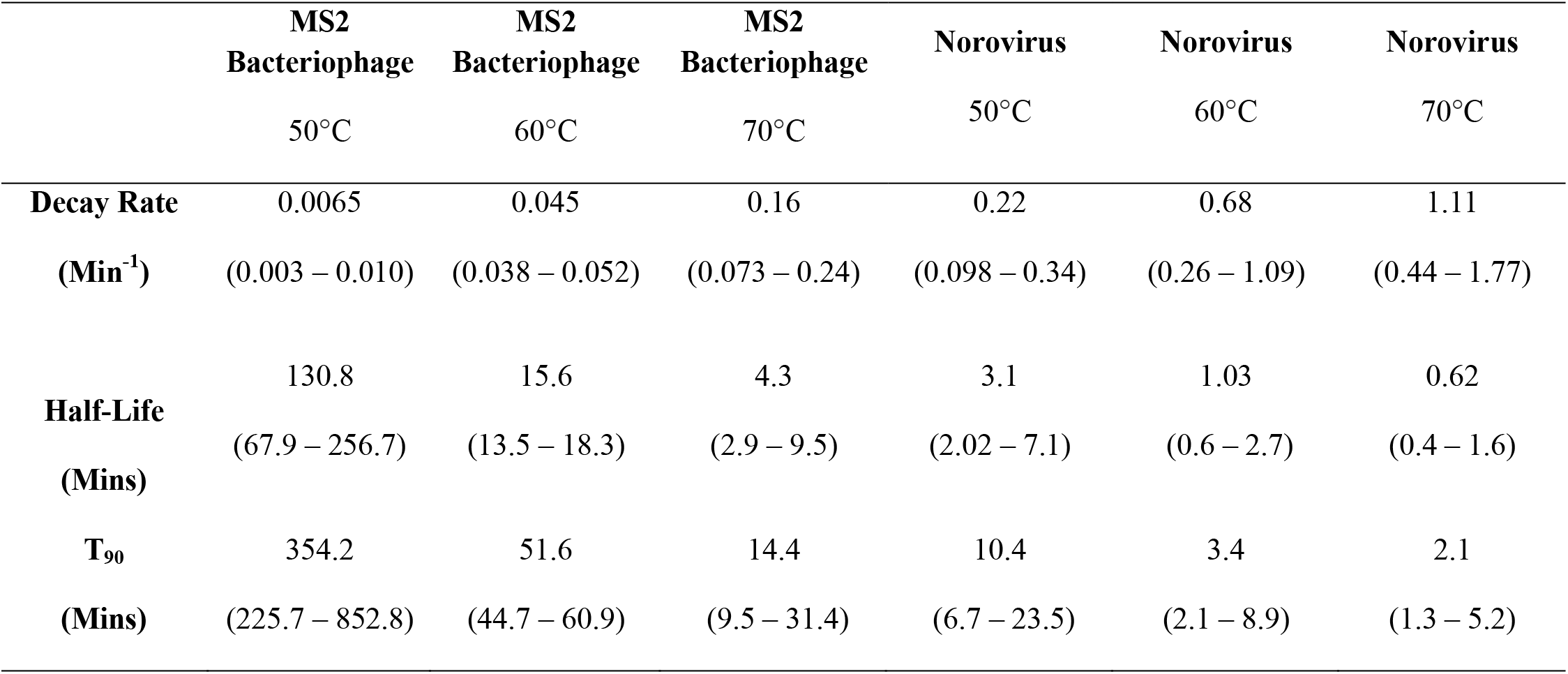
Summary of Decay Equations for Viable MS2 Bacteriophage and Norovirus. The values shown are the mean and the (95% CI).

|  | MS2<br>Bacteriophage<br>50°C | MS2<br>Bacteriophage<br>60°C | MS2<br>Bacteriophage<br>70°C | Norovirus<br>50°C | Norovirus<br>60°C | Norovirus<br>70°C |
| --- | --- | --- | --- | --- | --- | --- |
| **Decay Rate** | 0.0065 | 0.045 | 0.16 | 0.22 | 0.68 | 1.11 |
| **(Min<sup>-1</sup>)** | (0.003 – 0.010) | (0.038 – 0.052) | (0.073 – 0.24) | (0.098 – 0.34) | (0.26 – 1.09) | (0.44 – 1.77) |
| **Half-Life** | 130.8 | 15.6 | 4.3 | 3.1 | 1.03 | 0.62 |
| **(Mins)** | (67.9 – 256.7) | (13.5 – 18.3) | (2.9 – 9.5) | (2.02 – 7.1) | (0.6 – 2.7) | (0.4 – 1.6) |
| **T<sub>90</sub>** | 354.2 | 51.6 | 14.4 | 10.4 | 3.4 | 2.1 |
| **(Mins)** | (225.7 – 852.8) | (44.7 – 60.9) | (9.5 – 31.4) | (6.7 – 23.5) | (2.1 – 8.9) | (1.3 – 5.2) |

**Figure 1:**
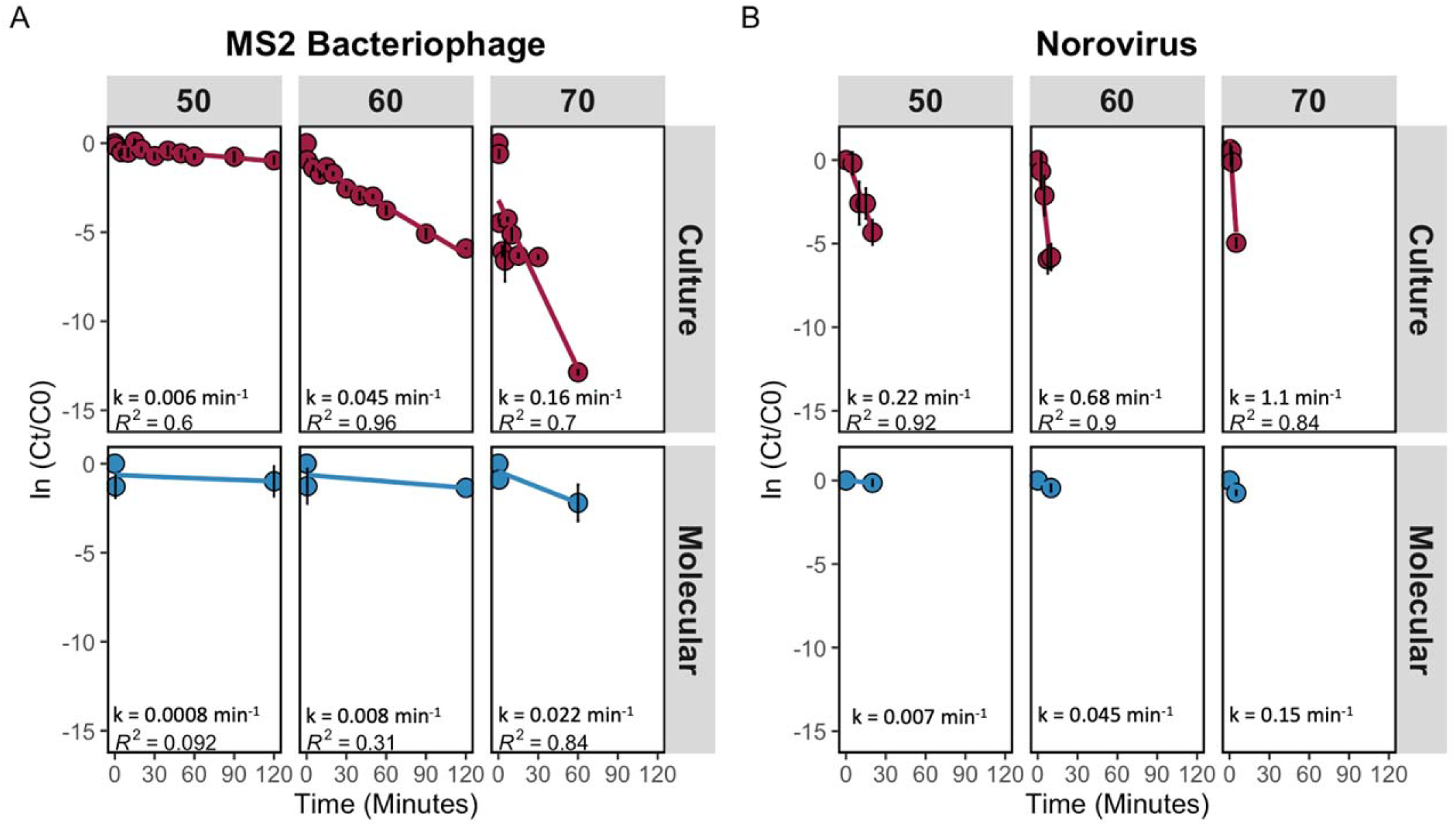
Monophasic Decay from Heat Inactivation. Decay rates and R-squared are shown on the plots. A) Monophasic decay of MS2 Bacteriophage with each column of panels representing a different experimental temperature. The top panels are from the culture assay, and the bottom panels are from RT-ddPCR. B) Monophasic decay of norovirus with each column of panels representing a different temperature and the rows showing the results from the HIE assay and the molecular detection of the input virus.

**Figure 2:**
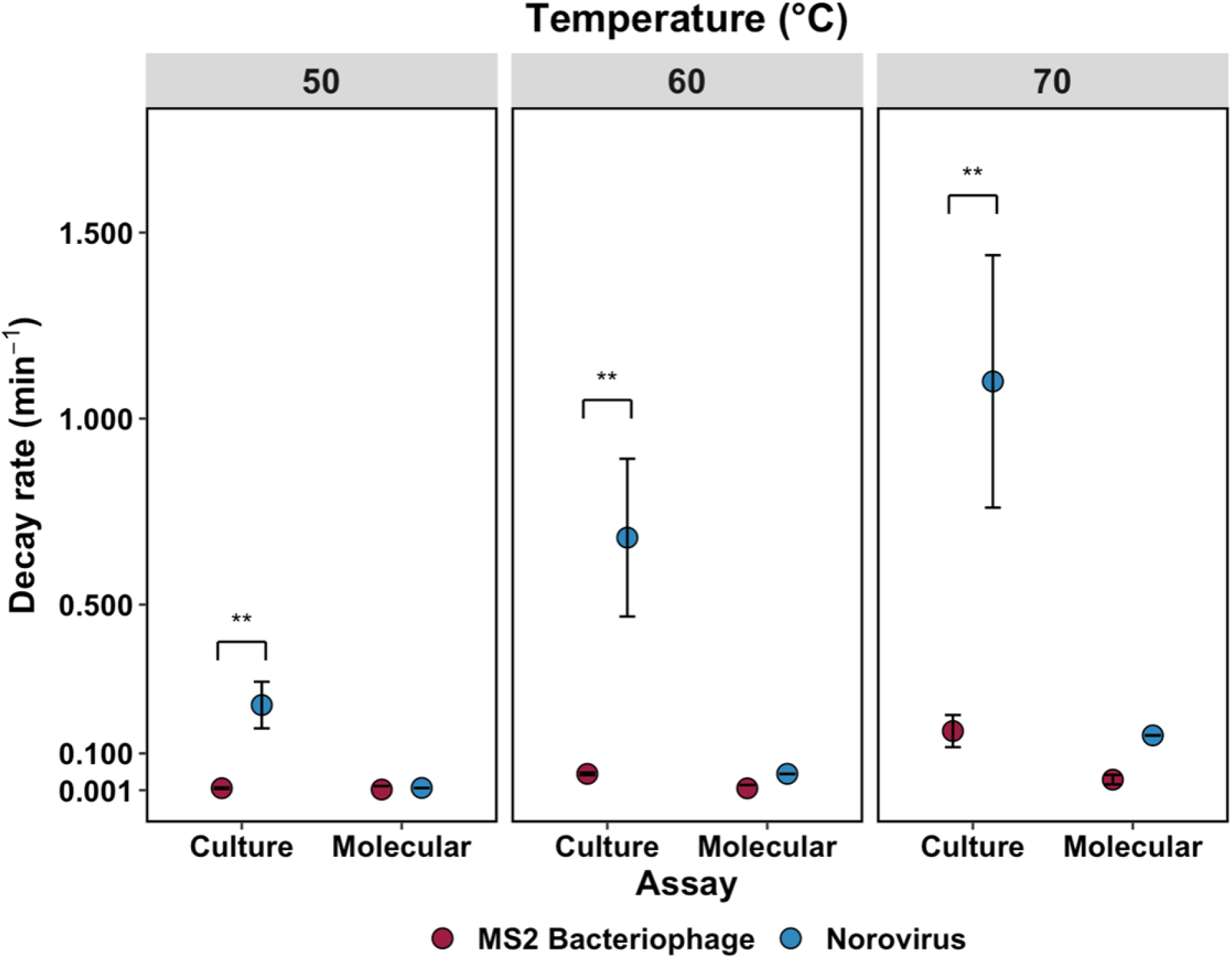
Decay Rate, k, versus temperature. The decay rates for each temperature for both viruses were plotted against temperature.

### Heat Inactivation of Norovirus

The monophasic decay of norovirus is shown in **Figure 1B**. The decay rate of viable human norovirus, shown by the slope of the line of best fit, increased as the temperature increased. The molecular data showed no significant change in the RNA signal with temperature. The decay rates for human norovirus were 0.22 min^-1^, 0.68 min^-1^, and 1.11 min^-1^ for 50°C, 60°C, and 70°C, respectively. Summaries of the monophasic decay rates, half-life, and T_90_ are shown in **Table 1**, and human norovirus decay rates were plotted against temperature, as shown in **Figure 2**; for each degree increase in temperature, the decay rate increased by 0.04 min^-1^.

## DISCUSSION

Heat is an effective treatment for the inactivation of enteric viruses in various matrices, including food (e.g., pasteurization) and wastewater (e.g., anaerobic digestion). Heat treatments are an effective and economical method for inactivating pathogens^4,16^ and have historically been used in the food industry, typically with low temperatures and long contact times to avoid product alterations. This study showed heat is a highly effective treatment method for the inactivation of human norovirus and shorter exposure times are necessary at higher temperatures to achieve the same level of norovirus inactivation.

MS2 bacteriophage, an icosahedral, single-stranded RNA virus, is a surrogate of interest for human norovirus.^17^ *Escherichia coli* is the host for MS2 bacteriophage; therefore, it can be detected in human feces and presents a similar seasonality with peaks in the winter months as norovirus.^18^ MS2 bacteriophage is commonly used as a surrogate virus for human norovirus because there is no requirement to have mammalian cell culture facilities, and the ease of culturing allows for relatively simple infectivity analyses.^17,19^ This phage is also non-pathogenic for humans, and high concentrations of phage particles, necessary for inactivation studies, are relatively easy to achieve.^18^ MS2 bacteriophage had significantly higher decay rates at each temperature than human norovirus, indicating less heat susceptibility for MS2 bacteriophage, as shown in **Figure 2**. MS2 bacteriophage is a conservative surrogate virus for human norovirus as inactivation occurs much faster with viable norovirus. Previous studies using RT-qPCR also determined MS2 bacteriophage to be a conservative surrogate virus; however, this study is the first to analyze viable norovirus instead of solely using the molecular signal for comparisons.^2^ Due to the cultivation methodologies, there were different matrices used for human norovirus and MS2 bacteriophage in this study and the impact of these matrices are not known and should be assessed in future work.

Experiments using other enteric viruses and human norovirus surrogates were conducted by Bozkurt et al., which analyzed the thermal inactivation of murine norovirus, feline calicivirus, and hepatitis A virus.^20,21^ In cell culture media, the T_90_ ranged from 19.95 - 20.23 min at 50°C and 0.56 - 0.94 min for 60°C for feline calicivirus, 34.49 - 36.28 at 50°C and 0.57 - 1.09 min for 60°C for murine norovirus, and were 56.22 min at 50°C and 2.67 min at 60°C for hepatitis A virus. Compared to the results of this study, MS2 bacteriophage has higher T_90_ than other human norovirus surrogates, indicating these surrogates using mammalian cells may be more indicative of the behavior of viable human norovirus.

Molecular methods are commonly used to determine the presence of viruses in samples; however, these methods cannot discern between viable and non-viable viruses. The results from these experiments show that there was no significant difference in the starting RNA signal and the RNA signal at the end of the experiment for both norovirus and MS2 bacteriophage, whereas there were significant differences in viable virus concentrations, as shown in **Figure 2**. A study conducted by Pecson et al. found an 8.6 log loss of MS2 bacteriophage at 72°C for 3 min; however, the qPCR signal loss was 1.6 log, further showing the discrepancy between loss of infectivity and loss of molecular signal.^19^ Overall, molecular methods largely underestimate treatment efficiency, especially when used to determine the efficacy of heat treatments.^22^

Viability pre-treatments, such as intercalating dyes and RNases, could eliminate free RNA and RNA from structurally compromised virions to further demonstrate the disconnect between molecular detection methods and viability. Further research is needed to determine optimal methods for these assays, as inhibition still results in the molecular signal not representing the viability of inactivated viruses.^7^

## Conclusions

This study shows heat is an effective treatment to inactivate human norovirus in an aqueous solution. Our data also show that human norovirus decay is significantly faster than MS2 bacteriophage decay, showing that MS2 bacteriophage is a conservative surrogate for human norovirus under heat inactivation. Finally, our data demonstrate that molecular measurements do not correlate with viable human norovirus under heat inactivation.

